# Flexible coding of memory and space in the primate hippocampus during virtual navigation

**DOI:** 10.1101/295774

**Authors:** Roberto A. Gulli, Lyndon Duong, Ben Corrigan, Guillaume Doucet, Sylvain Williams, Stefano Fusi, Julio C. Martinez-Trujillo

## Abstract

Hippocampal maps of space change across tasks. The mechanisms of this effect remain unclear. To examine this, we recorded activity of hippocampal neurons in monkeys navigating the same virtual maze during two different tasks: a foraging task requiring only cue guided navigation, and a memory task also requiring context-object association. Within each task, individual neurons had spatially-selective response fields, enabling a linear classifier to decode position in the virtual environment in each task. However, the population code did not generalize across tasks. This was due to sensory and mnemonic coding of non-spatial features and their associations by single neurons during each period of the associative memory task. Thus, sensory and mnemonic representations of non-spatial features shape maps of space in the primate hippocampus during virtual navigation. This may reflect a fundamental role of the hippocampus in compressing information from a variety of sources for efficient memory storage.

## Introduction

The hippocampus is a phylogenetically ancient structure that receives highly processed input from all sensory modalities (Felleman and Van Essen, 1991; van den Heuvel and Sporns, 2011). Processing of varied sensory inputs in the hippocampus serves two complex cognitive functions: memory and spatial navigation. Evidence for hippocampal involvement in both of these functions spans decades, though each is supported by largely disparate bodies of literature (Buzsáki and Moser, 2013; Eichenbaum and N. J. Cohen, 2014; Ekstrom and Ranganath, 2017; Schiller et al., 2015).

Neurophysiological evidence for hippocampal involvement in memory recall began with Penfield’s evocation of rich and vivid memories after direct electrical stimulation in epileptic patients in the 1930s. Stimulation of the medial temporal lobe, including but not limited to the hippocampus evoked “experiential hallucinations”, in which patients would recall remote episodes of their lives in rich detail: a “reactivation of a strip of the record of the stream of consciousness” (Penfield, 1958). These studies were complemented by contemporary observations that bilateral ablation of the hippocampus led to profound memory deficits (Milner and Penfield, 1955; Penfield and Milner, 1958; Scoville, 1954; Scoville and Milner, 1957). Immediately, efforts were made to define a non-human primate model that clearly replicated these deficits to understand the neurophysiology that was disrupted in patients. However, results were equivocal (Correll and Scoville, 1965a; 1965b; Orbach et al., 1960); only decades after the description of HM, analogous deficits in memory were observed in humans and non-human primates after ablation of the hippocampus, amygdala and rhinal cortex (Mishkin, 1978). At that time, it became clear that associative, relational, and/or contextual information relevant to memory was encoded by the hippocampus, providing a parsimonious explanation of the memory deficits observed after hippocampal perturbation.

In the time between these behavioral findings, observation of single hippocampal neuron activity in rodents during a wide variety of behaviours became commonplace. This culminated in a succinct description of single neurons that fire action potentials when rodents occupied a particular region of a familiar environment (O’Keefe and Dostrovsky, 1971). Early studies showed place cells that fired in a direction- and behaviour-invariant manner, suggesting the hippocampus encodes a complete allocentric cognitive map of the environment for spatial navigation (O’Keefe and Nadel, 1978). The discovery of place cells ushered in a new era of hippocampal research on spatial responses of hippocampal neurons and ensembles (Moser et al., 2017). An emergent consensus from this body of research is that spatial representations in the hippocampus can be modulated by a variety of environmental, cognitive, and behavioral factors (Fenton et al., 2010; Gothard et al., 2001; Markus et al., 1995). Relatively few studies have extended these observations to non-human primates, which typically report complex spatial response fields in hippocampal neurons that are modulated by direction, spatial view, motivational goals, and “action context” (Miller et al., 2013; Rolls and Xiang, 2006; Wirth et al., 2017).

Extensions to the cognitive mapping theory suggest the hippocampus more generally builds maps of low-dimensional features along a continuous scale, including time, sound, and other abstract concepts (Aronov et al., 2017; Schiller et al., 2015). Recently, however, it has been inversely proposed that the role of the hippocampus in mapping these dimensions is related to its fundamental role in memory (Eichenbaum, 2017a; Hardcastle et al., 2017; Stachenfeld et al., 2017). Direct comparisons of how continuous spatial maps are modulated by discrete sensory and mnemonic representations in hippocampal neurons are scarce for several reasons. First, a large methodological gap exists between studies of mapping in rodents and discrete tests of memory performance in primates (Buzsáki and Moser, 2013; Eichenbaum and N. J. Cohen, 2014; Ekstrom and Ranganath, 2017; Schiller et al., 2015). Second, diverse behavioural repertoires across species based on the structural organization of sensory systems and corresponding reorganization of sensory inputs to the hippocampus make cross-species comparisons of activity amongst hippocampal neurons complex (Preuss, 2000).

To directly examine factors that may influence the stability and expression of cognitive spatial maps, we recorded single neuron activity from the hippocampi of two male non-human primates (*Macaca mulatta*). Electrode trajectories targeted the superior aspect of the right mid-posterior hippocampus (predominantly CA3). Trajectories were planned and verified during electrode insertion using MRI-guided neuro-navigation (**Figure 1A** & **B**, **Figure S1** & **S2**). Within each recording session, the monkeys completed three different tasks during simultaneous recording of eye movements and single-neuron activity from the hippocampus.

**Figure 1.**
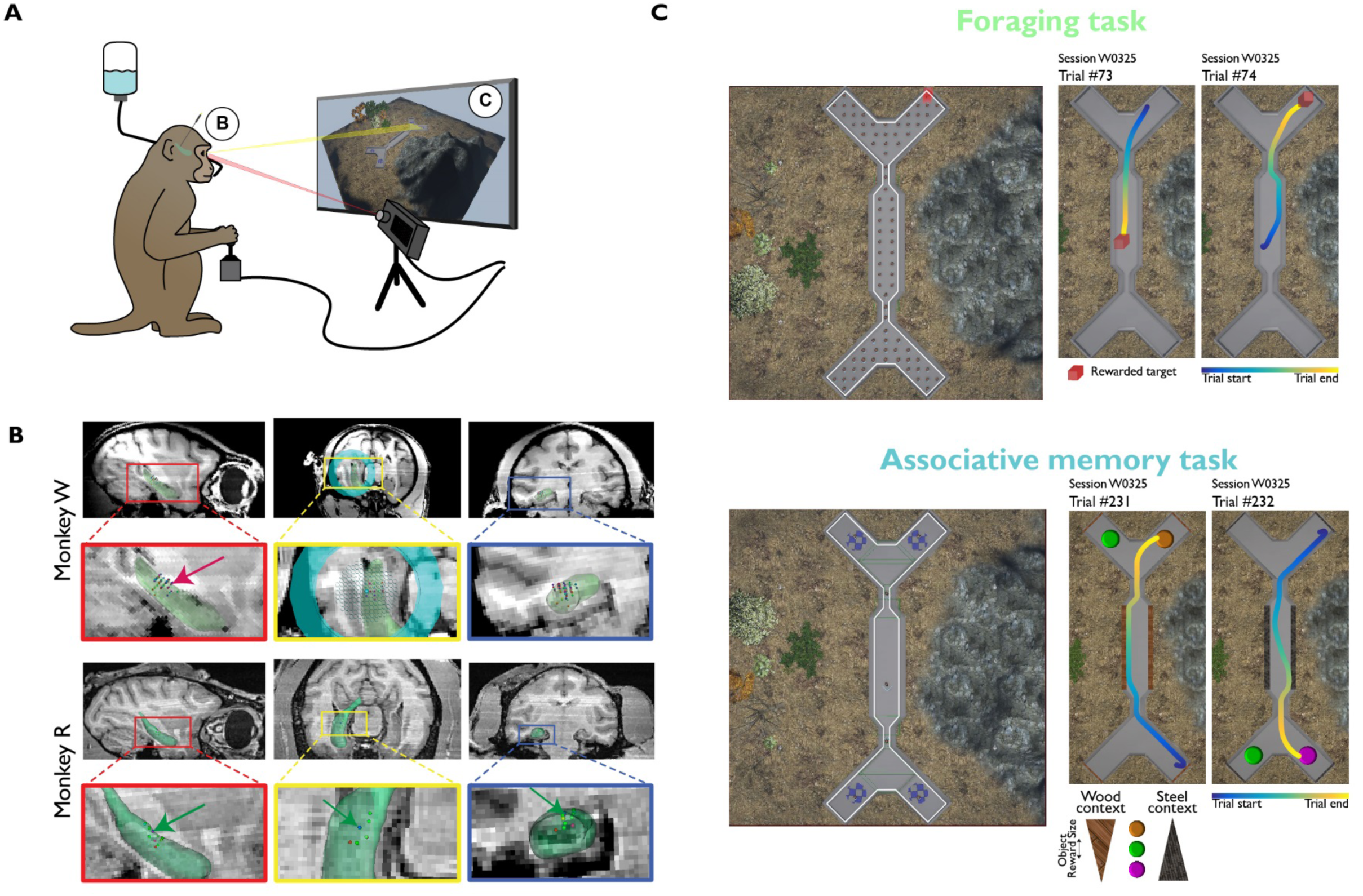
Tasks and recording locations. **(A)** Monkeys were seated in front of a computer monitor and used a two-axis joystick to navigate through virtual reality X-Maze and complete two different tasks **(B)** Single-neuron recording locations from the right hippocampi (green reconstruction) of two monkeys. Arrows mark the recording locations of example neurons shown in subsequent figures **(C)** Overhead view of potentially rewarded locations and example trial trajectories in the X-Maze during the Foraging (top), and context-object Associative Memory task (bottom) Related to **Figures S1**, **S2**, and **S3**.

Two tasks – the foraging task and the associative memory task – required free navigation in the same custom-built virtual reality environment (X-Maze) using a two axis joystick (Doucet et al., 2016) (**Figure 1A** & **C**). The spatial layout of the environment was unchanged across tasks.

In the foraging task, animals navigated through the X-Maze towards a red volume to receive a juice reward. The red volume was randomly assigned to one of 84 locations in the maze, and randomly repositioned at a different location every time the subject reached it (**Figure 1C**, top; **Movie S1**; https://youtu.be/aWvheMzxMJo).

In the associative memory task, monkeys learned a reversed two-context, three-object reward value hierarchy embedded into the X-Maze (**Figure 1C**, bottom; **Figure S3**; **Movie S2**; https://youtu.be/RHx9Lw65oDw). Within a trial, context was cued by applying a wood or steel texture to the maze walls when the subject reached the central corridor (**Figure S3 B** & **C**, position a). When subjects reached the branched point opposite where they started (**Figure S3 B** & **C**, position b), one of the three possible colored discs appeared in each arm of the maze. The context and object pair on every trial was selected randomly. In every session, a new set of three colored discs was used with the same two contexts. Thus, a new conditional association between context and objects was learned daily.

In each session, the VR tasks were bookended by a cued saccade task wherein monkeys were rewarded for making saccades to small white dots on a gray screen. This task was used to calibrate and validate accuracy of eye-on-screen position and examine whether hippocampal neurons exhibit screen position or saccade direction selectivity.

## Results

### Single neuron firing rates are tuned for spatial position in both tasks

To characterize place-selectivity in individual neurons, the virtual environment was spatially binned into an isometric two-dimensional pixel grid covering the entire maze. In each task, occupancy time and number of spikes observed were used to compute empirical per-pixel firing rates for each neuron. A wide variety of spatial distributions of spikes could be seen across neurons and across tasks (**Figure 2A** & **B**); changes in spatial distribution of activity were not due to changes in neuronal isolation (**Figure S2**). Across all neurons, firing rates were higher in the branching points and arms of the maze during the associative memory task than during the foraging task (**Figure 2C**).

**Figure 2.**
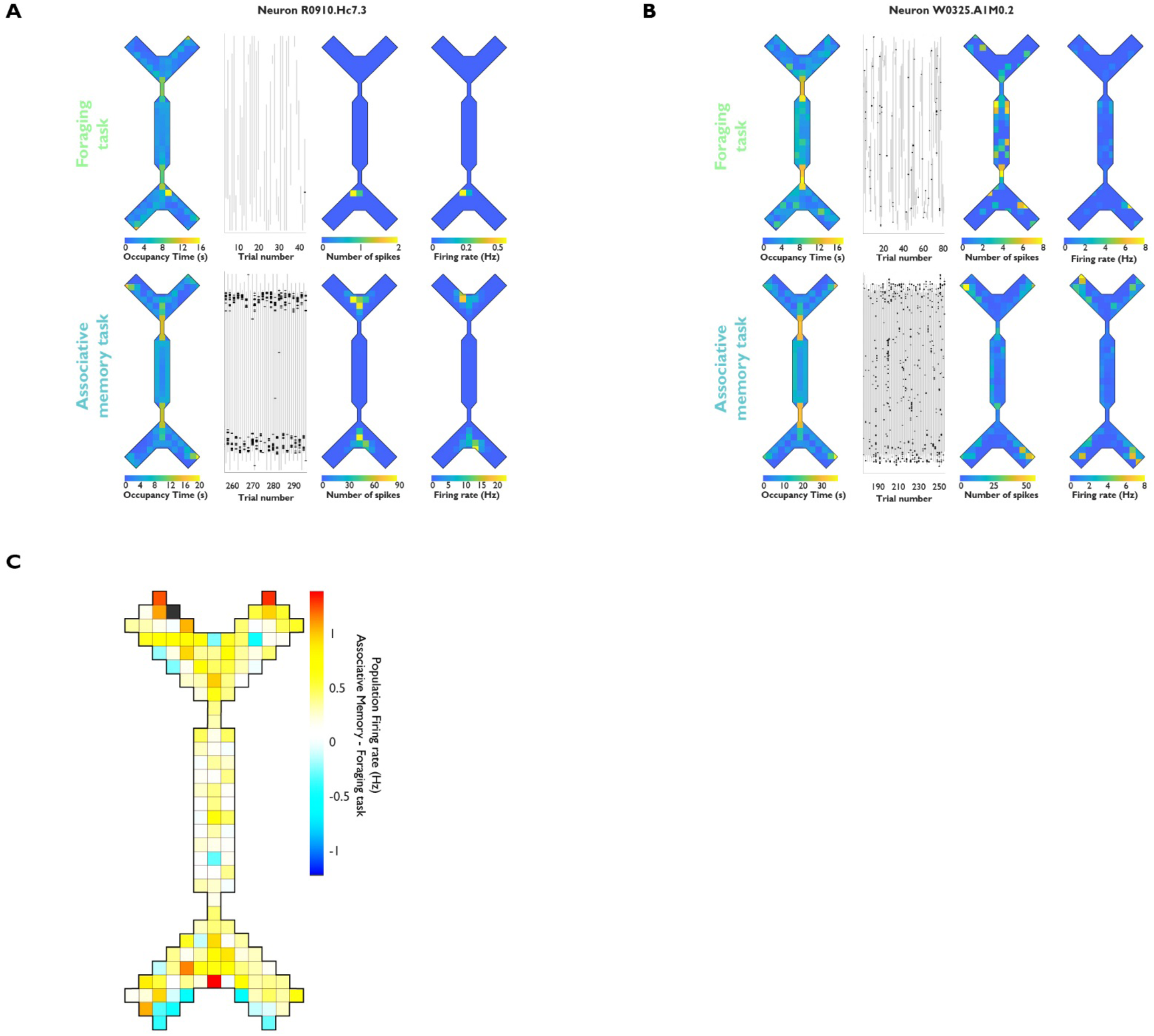
Example neuron firing rates during the Foraging and Associative Memory task. **(A)** Example neuron R0910.Hc7.3. Top row shows data collected during the Foraging task; bottom row shows data collected during the Associative Memory task. Spatial map of total time spent in each map pixel (left/first column), trial-by-trial spike locations along the north-south dimension (second column), number of spikes in each pixel (third column), and per-pixel firing rate (right/fourth column) **(B)** Example neuron W0325.A1M0.2. Conventions are the same as in (A) **(C)** Average cross-task firing rate difference for all neurons

To determine which pixels’ empirical firing rates were higher than expected by chance, spatial position (pixel number) and neural activity were vectorized with 1ms resolution in each task. The neural activity vector was circularly shifted to change the spike location on each trial 1000 times, creating a permutation-derived null distribution of firing rates for each pixel, each neuron, and each task. The number of neurons with firing rates that are statistically elevated (alpha=0.05/number of occupied pixels for that neuron; see Methods) in each pixel and in each task can be seen in **Figure 3A**. The X-Maze was then divided into 9 similarly-sized areas for further analyses (4 maze arm areas, 2 branch areas, 3 corridor areas). Spatial response fields for each neuron were defined as any of the 9 maze areas with statistically elevated firing in one or more pixels.

**Figure 3.**
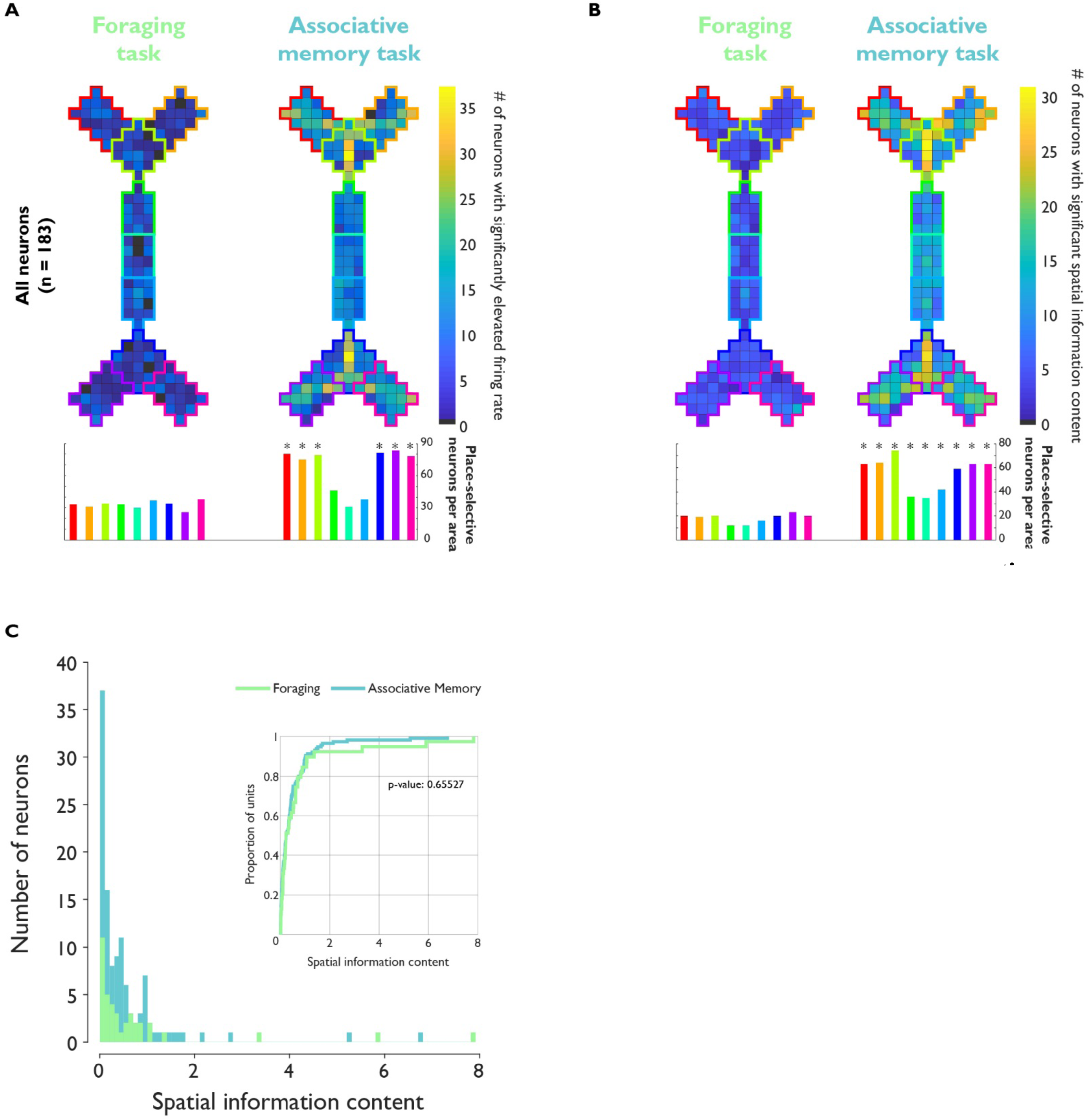
Firing rate and Spatial Information Content in both tasks. **(A)** Spatial histogram showing the number of neurons with statistically elevated firing rate in each pixel in both tasks (top). The summarized histogram (bottom) shows the number of neurons with at least one significant pixel in each maze area. *significantly different proportion across tasks; McNemar’s test of equal proportions, p<0.05, Bonferroni-corrected **(B)** Histograms showing the number of neurons with statistically elevated spatial information content. Conventions are the same as in (A) **(C)** Histogram of significant spatial information content values in each task. Only neurons with empirical spatial information values above the 95^th^ percentile of circularly shifted permutation test values are included. Inset: Cumulative distribution of spatial information content in each task Related to **Figures S4** and **S5**.

In the foraging task, 55.7% of neurons had spatial response fields in at least one maze area (102/183 neurons, median 1; **Figure S4A**, green bars). These were homogeneously distributed throughout the maze (p=0.91, *χ*^2^(8) = 3.29; **Figure 3A**, left column). In the associative memory task, 70.0% of neurons also had spatial response fields. In contrast with the foraging task, these were not homogeneously distributed throughout the X-Maze (p<1⨉10^−7^, *χ*^2^(8) = 52.3; **Figure 3A**, right column). Of all neurons with spatial response fields in both tasks (49.7%; 91/183), neurons were likely to gain fields in the arms and branching points of the maze during the associative memory task compared to the foraging task (**Figure S4B**).

One possibility is that place-specific firing can be ascribed to selectivity for eye-on-screen position (Killian et al., 2012) coupled with biased gaze behavior within or across tasks. We compared saccade direction and gaze position selectivity in all three tasks for neurons with sufficient numbers of saccades and fixations in all tasks (n = 92 neurons; Methods). In the cued saccade task, foraging task, and associative memory task, 7.6%, 41.3%, and 42.4% of neurons were selective for saccade direction, respectively. In these three tasks 28.2%, 43.4%, and 72.8% of neurons were selective for gaze position. Critically though, no neurons were selective for the same saccade direction across tasks, and only 2 neurons (2.2%) were selective for at least one gaze position across all tasks. Thus, saccade direction and gaze position invariably affect only a small proportion of hippocampal neurons and cannot explain the dramatic changes in spatial selectivity across tasks.

### Single neuron spatial information content varies across tasks

An additional measure of spatial encoding in individual neurons was used to quantify firing specificity in each task; spatial information content (SIC) quantifies how many bits of information about the location of the subject are transmitted per action potential (Markus et al., 1994; Ravassard et al., 2013; Skaggs et al., 1993). Empirical and circularly shuffled control SIC values were computed using occupancy vectors and firing rate vectors, much the same as previously described (see Methods). **Figure 3B** shows the number of neurons with SIC values that exceed the statistical threshold for significance compared to the null distribution of each pixel (alpha=0.05/number of occupied pixels).

In the foraging task, 27.3% of neurons had significant SIC summed over the entire spatial map. The location of pixels with significant SIC were homogeneously distributed throughout the maze (p=0.59, *χ*^2^(8) = 6.56; **Figure 3B**, left column). In the associative memory task, 67.2% of neurons showed significant SIC. The location of pixels with significant SIC were not homogeneously distributed throughout the maze (p=0.0004, *χ*^2^(8) = 28.5; **Figure 3B**, right column), and the proportion of neurons with significant SIC during the associative memory task exceeded that observed in the foraging task for all maze areas (McNemar’s test for equal proportions; p-value range 0.005 - 1×10^−7^). Though more neurons exhibited significant SIC in the associative memory task than the foraging task, the distribution of empirical SIC values was not different between tasks (**Figure 3C** and **Figure S5**; two-sample Kolmogorov Smirnoff test, p=0.66).

### Spatial position can be decoded from population activity

The previous results show that individual neurons contain information about space in each task, but statistical descriptions of selectivity in single neurons may not be sufficient in capturing the wealth of information encoded by neural populations (Fusi et al., 2016; Rigotti et al., 2013). The stability of a holistic population code for space and the optimal spatial reference frame cannot be assessed in single neurons. Therefore, we used the firing rates from the entire population of neurons to decode subject position in the maze within and across tasks, and using allocentric and direction-dependent spatial reference frames (**Figure 4A**).

**Figure 4.**
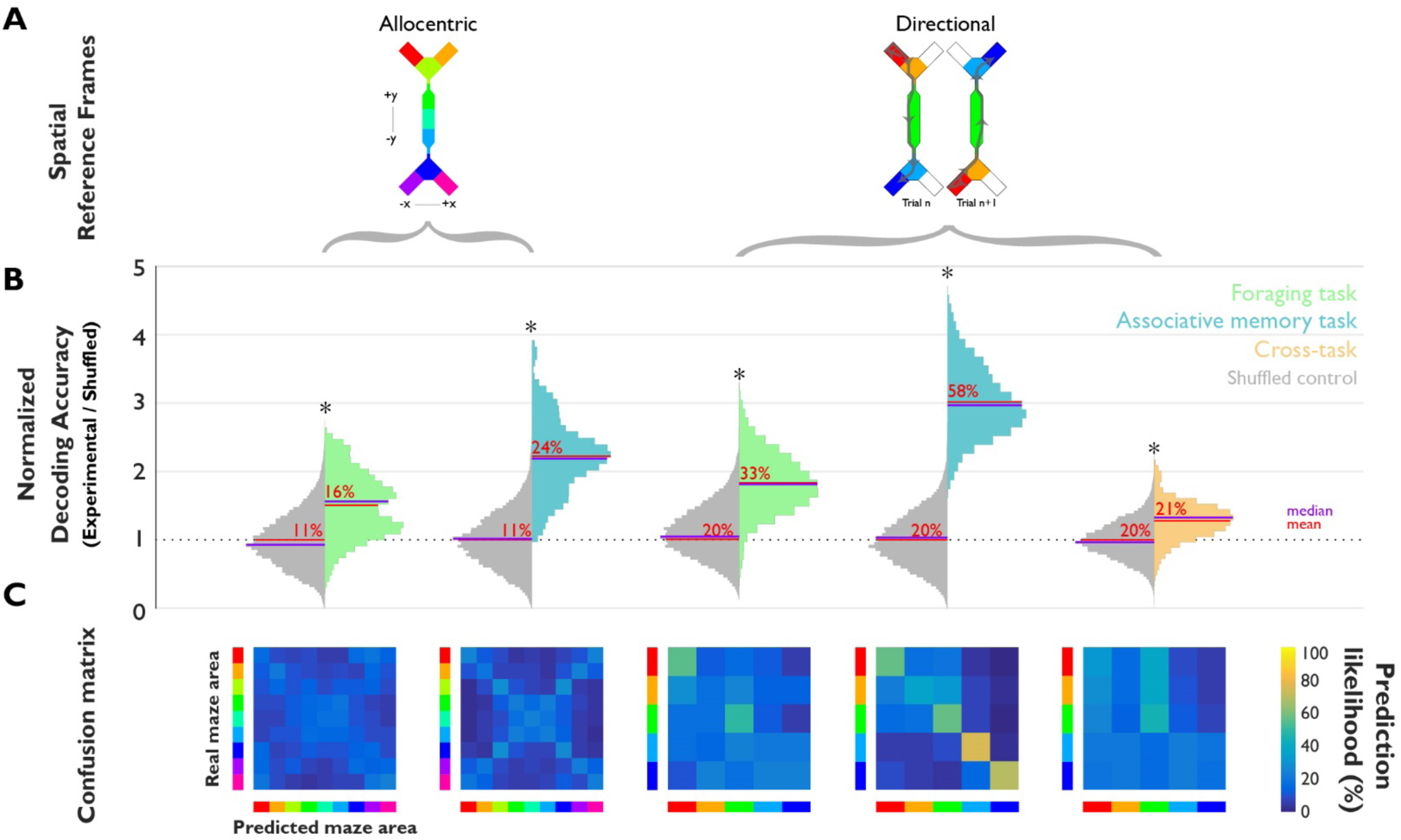
Hippocampal ensemble prediction of spatial position with a linear classifier. **(A)** Spatial reference frames used in each of the lower panels of spatial classification analyses **(B)** Distributions of classification accuracies for spatial position in the foraging (green) and associative memory (blue) tasks as well as cross-task testing and validation (orange). *p<0.05 real vs. shuffled distribution (grey distributions), Wilcoxon ranksum. Red bars and numbers, mean; purple bars, median **(C)** Confusion matrices for each classification analysis. Real location=rows; predicted location=columns; colormap, prediction likelihood

We used a multi-class support vector machine with a linear kernel to decode the subject’s position in space using neuronal firing rates (Fan et al., 2008; Methods). Briefly, an ensemble was constructed using firing rates from each neuron in 10 passes through each maze area for each task; neurons with less than 10 trials in any condition were excluded (n=152 neurons included). Decoding accuracy was tested using cross-validation. The ensemble was stratified into 5 folds, with 4 of these used for parameter normalization and model training, and the remaining fold used for model testing. A null distribution of decoding accuracies was derived by permuting the labels of the testing fold. This procedure was repeated 100 times, yielding distributions of empirical (**Figure 4B**, colored distributions) and null (**Figure 4B**, grey distributions) decoding accuracy. Statistical differences between accuracy distributions were assessed via Wilcoxon ranksum.

The classifier predicted position in the maze above chance levels in both tasks when using an allocentric spatial reference frame (**Figure 4B**, first and second column; foraging: 1.50±0.45 times permuted control (mean±SD); p<10^−12^; associative memory: 2.22±0.53 times permuted control; p<10^−31^). However, accuracy was poor (foraging task Cohen’s *κ*=0.06; associative memory task Cohen’s *κ*=0.15; Methods). The classifier systematically confounded structurally similar areas of the maze in both tasks, as evidenced by the X-shaped distribution of predictions in the allocentric decoder confusion matrices (**Figure 4C**).

Decoding position using an idiothetic, direction-dependent reference frame, prediction accuracy was 1.71±0.39 (p=0.09 compared to allocentric; **Figure 4B**; third column). In the associative memory task, decoding accuracy improved to 3.01±0.48 (p<10^−10^ compared to allocentric; **Figure 4B**, fourth column). These results indicate that populations of hippocampal neurons encode spatial position more reliably in a direction-dependent idiothetic reference frame, consistent with previous findings across species (Acharya et al., 2016).

### Cross-task comparison of population models of spatial position

Though subjects’ position in space could be decoded from the population of neurons in each task, it is not clear whether coding of space generalized across tasks; that is, whether an abstract representation exists, invariant to the task that the animal is engaged in (Saez et al., 2015). To determine whether the cognitive map of space generalized across tasks, we trained a linear support vector machine using trials from the foraging task and tested using trials from the associative memory task. Classification accuracy fell near chance (**Figure 4**, orange, rightmost column; 1.28±0.25 times permuted control accuracy; p<10^−11^; *κ*=0.08). Results were unchanged when the training and testing sets were swapped. This indicates that coding of spatial position by the ensemble of recorded neurons does not generalize across tasks and suggests that the spatial information carried by the population of recorded neurons changes across tasks.

The poor abstraction of spatial representation across tasks could be due to a global change across the entire X-Maze, or differences in parts of the maze where behavioral demands differed across tasks (branches and arms). At these points, object- and context-driven comparison and selection of targets is required only in the associative memory task. On the other hand, cue guided navigation through the central corridor was a common requirement across tasks. By examining the confusion matrix for the cross-task decoding analyses, it is clear that cross-task predictions were most accurate in the corridor of the maze (**Figure 4C**, farthest right).

### Individual neurons encode sensory and mnemonic features of the associative memory task

The previous results show where in the X-Maze the population code for space changes, but not which parameters of the task are being encoded in each area. Changes in the population model across tasks may be due to neurons encoding features of the associative memory task (context, object color, and their conjunction) at the locations in the maze where they become task relevant. To examine this encoding, we regressed firing rate for each neuron against the trial-varying parameters of the associative memory task in each period. This was done using the context and object of a given trial and the firing rate in each period of the same trial (sensory encoding; **Figure 5**, left plots), or the firing rate in each period of the next trial (memory encoding; **Figure 5**, right plots).

**Figure 5.**
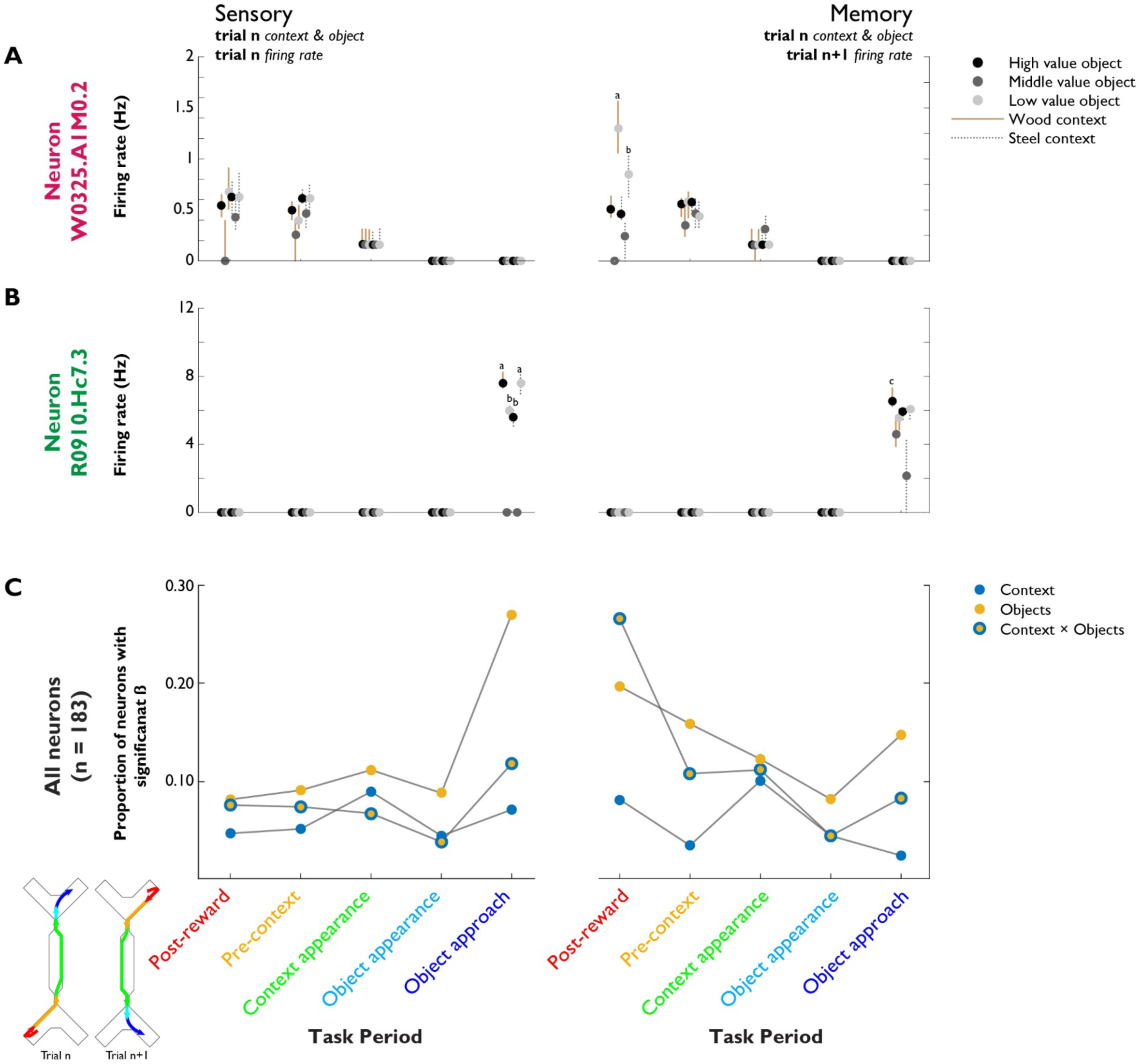
Non-spatial feature selectivity in the associative memory task. Selectivity of each neuron for non-spatial features of the associative memory task. Selectivity is computed using trial parameters of trial n and firing rates from the same trial (sensory; left plots), or trial n+1 (memory; right plots). **(A)** Example neuron W0325.A1M0.2. an object-value selective neuron. Letters denote categories with significantly different firing rates within that condition (p<0.05, Bonferroni adjusted) **(B)** Example neuron R0910.Hc7.3, an object-color selective neuron. Conventions are the same as in (A) **(C)** Proportion of single neurons selective for non-spatial associative memory task features: context (blue dot), chosen object color (yellow dot), and the combination of these two (yellow dot, blue outline) Related to **Figure S6**.

The example neuron shown in **Figure 2B** fired most in the arms of the maze. Plotted as a function of trial period in the associative memory task, this neuron fires immediately after the reward was delivered; post-reward firing was not observed in the foraging task (**Figure S7**). This example neuron was selective for previous trial features during the post-reward period (**Figure 5A**); firing rate was highest following trials where the lowest value object was chosen, and no reward was delivered (p<10^−4^, Kruskal Wallis). This encoding cannot be explained by sensory features such as object color, context or reward size alone; but only by the conjunction of these features, even though none of these features were visible during the post-reward period.

The example neuron shown in **Figure 2A** fired exclusively between the subject’s initial turn towards the chosen object and the moment of first contact (goal approach period; **Figure S7**). This neuron was most active during approaches to a single object color regardless of context, and did not fire when approaching an intermediate value object (**Figure 5B**; p<10^−20^, Kruskal Wallis).

On a population level, a diverse array of selectivity for associative memory trial features was observed. The chosen object color was most robustly encoded during the object approach (46/183 neurons, 25.14%; **Figure 5C**, left, yellow dots). The conjunction of object and context of the previous trial was encoded by the same number of neurons during the post-reward period (**Figure 5C**, right, yellow dots with blue outline). Encoding of trial context peaked in the corridor of the maze (sensory: 16/183 neurons, 8.7%, memory: 18/183 neurons, 9.8%; **Figure 5C**, blue dots).

These results show non-spatial sensory and mnemonic encoding changes across trial periods of the associative memory task. It is not known whether these functions are supported by a common or separate population of hippocampal neurons. To compare sensory and memory encoding of each neuron for each trial parameter, we correlated F-statistics from each neuron’s encoding model during goal approach (sensory) and post-reward (memory) periods (**Figure 6**). Similar proportions of neurons showed prospective and retrospective coding of contexts (**Figure 6A**, left, p=0.68, McNemar’s test of equal proportions) or objects (**Figure 6A**, middle, p=0.12, McNemar’s test of equal proportions). However, for the combination of object and context, the proportion of neurons showing memory coding was significantly larger than sensory coding (**Figure 6**, left, p=0.0005, McNemar’s test of proportions). The strength of sensory versus memory coding in individual neurons was not correlated for trial context (**Figure 6B**, Spearman rho 95% confidence interval = −0.12–0.25); in contrast, sensory and memory coding for objects and combinations of contexts and objects were correlated (**Figure 6B** objects: Spearman rho 95% confidence interval = 0.13–0.48; context×object: 0.05–0.40).

**Figure 6.**
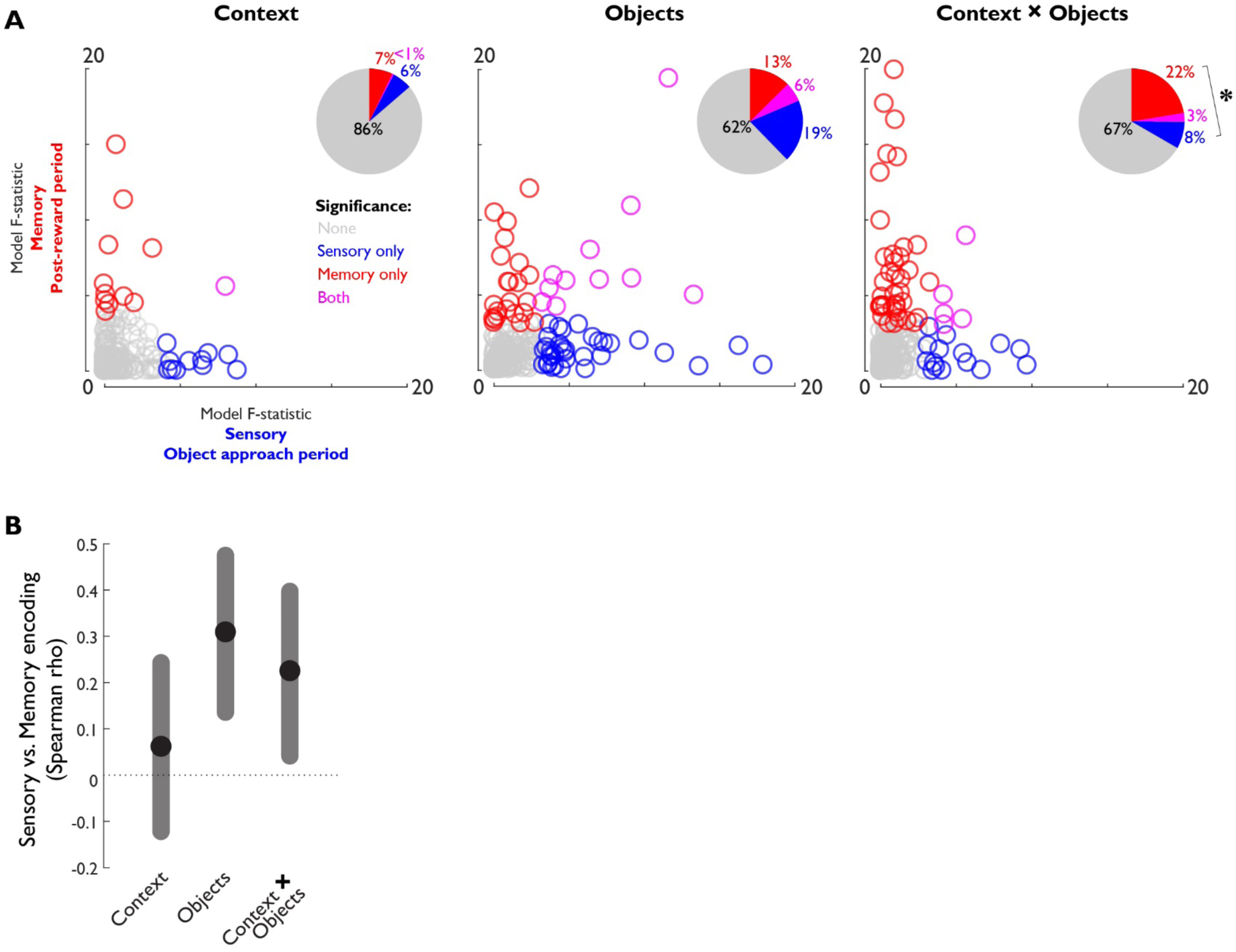
Sensory versus Memory encoding of trial features. **(A)** Sensory versus memory encoding model: fit parameter F-statistic comparison. Inset: percentage of neurons with significant fit parameters for each encoding model **(B)** Spearman correlation coefficient of encoding model F-statistics for each neuron between the foraging and associative memory task. Circle, median; shaded bars, 99% confidence interval

### Associative memory trial type is encoded by the population

Finally, we used a linear classifier to determine the accuracy with which the population of hippocampal neurons can predict the context and object pair of a given trial (associative memory trial type). One classifier was trained to predict context and object of the current trial from three sensory trial periods (context appearance, object appearance, and object approach; **Figure 7**, Sensory); another was trained using two memory trial periods (post-reward and pre-context; **Figure 7**, Memory). In both classifiers, each neuron’s firing rate in each trial period was used as an independent predictor. We used a logistic classifier with elastic net regularization to avoid problems of overfitting associated with having many predictors and a limited number of model training examples (Friedman et al., 2010).

**Figure 7.**
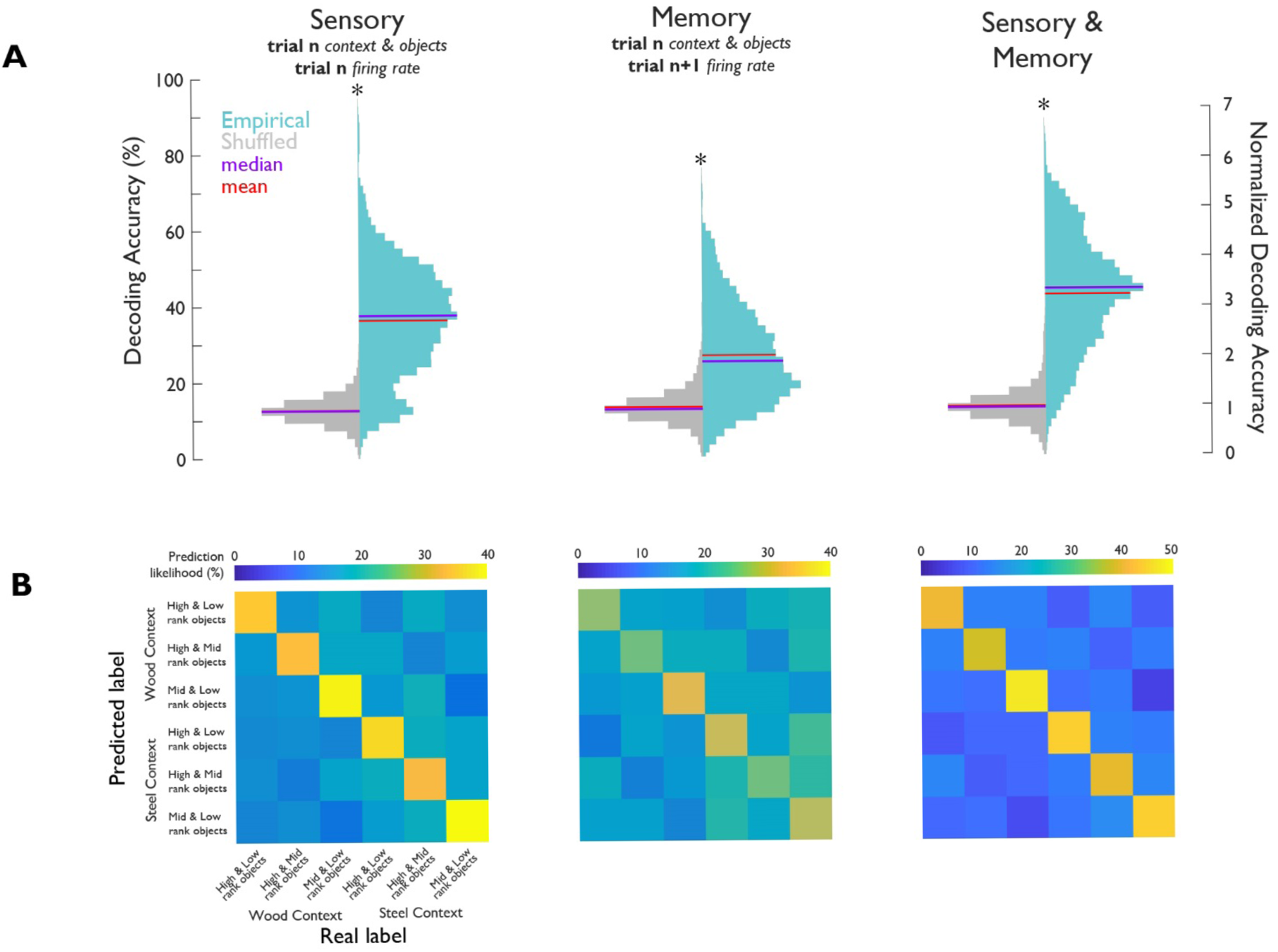
Trial type decoding in the associative memory task. **(A)** Distribution of classification accuracies from decoding analysis of trial type (trial context and object pair) in the associative memory task. *p<0.05 real vs. shuffled distribution (grey distributions), Wilcoxon ranksum. Red bars, mean; purple bars, median **(B)** Confusion matrix derived from the classification analysis. Predicted trial type=rows; real trial type=columns; colormap, prediction likelihood

Using only the sensory or memory trial periods, prediction accuracy was above chance (sensory: 2.88±1.13 times permuted control; p<10^−86^, Wilcoxon ranksum, Cohen’s *κ*=0.24; memory: 2.04±0.95; p<10^−58^ times permuted control, Wilcoxon rank-sum, Cohen’s *κ*=0.12). Using the firing rates from both the sensory and memory trial periods in a single classifier, classification accuracy further increased (3.29±1.12 times permuted control; p<10^−97^, Wilcoxon ranksum; Cohen’s *κ*=0.31).

## Discussion

In the current study, we recorded the activity of individual hippocampal neurons while monkeys performed a navigation task and an associative memory task in the same freely navigable maze. During both tasks, hippocampal neurons encoded spatial information, though single-neuron and population coding of space changed across tasks. Critically, observed changes to spatial coding across tasks were attributed to selectivity for trial-varying features of the associative memory task visible in the environment (sensory representation), and when they were no longer visible (mnemonic representation). These results extend previous observations of hippocampal encoding, showing that dynamic encoding of space can be attributed to sensory and mnemonic representation of perceptually and cognitively defined features.

### Spatial remapping in hippocampal neurons

Changes in single neuron and population coding for space in the current study may be interpreted as place cell remapping, as seen in studies of rodent hippocampal activity in real (Muller and Kubie, 1987) or virtual environments (Acharya et al., 2016). Several types of remapping have been defined: global, partial, local, and rate (Knierim and McNaughton, 2001; Moser et al., 2017). Global remapping occurs when all neurons with place-specific firing rearrange their preferred firing location. Partial remapping occurs when some, but not all recorded place cells change their preferred firing location in response to a global change in the environment. Local remapping occurs when some, but not all recorded neurons change their preferred firing location in response to a localized change in the environment (e.g. addition, removal, or movement of an object or barrier). In contrast, rate remapping occurs when a neuron’s preferred firing location is preserved, albeit at a significantly different rate (Leutgeb et al., 2005). The specific environmental or cognitive factors, and related thresholds that lead to each type of remapping are unclear (Moser et al., 2017).

In the current study, neither global nor rate remapping sufficiently captures the nature of cross-task changes in spatial and non-spatial encoding. Similarity in cross-task models for space in the central corridor of the maze, as illustrated in the cross-task confusion matrix in **Figure 4C**, suggests that not all neurons with place-specific activity change their contribution to spatial decoding across tasks. Since firing rates were scaled within tasks for these analyses, rate remapping could not explain reduced prediction accuracy in cross-task spatial decoding analyses. The localized change in cross-task models of space in the arms and branches of the maze are most akin to local remapping. Beyond this, we show that elevated firing of single neurons is attributed to sensory and mnemonic selectivity for specific features of objects at these locations in the associative memory task. The high proportion of neurons with sensory and mnemonic selectivity for non-spatial trial-varying features of the associative memory task suggest that it is the encoding of these features – rather than remapped selectivity for space per se – that explains changes in spatial representation across tasks. The nuances in spatial and non-spatial mnemonic encoding we have reported here have not been previously observed in the primate hippocampus.

### Non-spatial encoding in hippocampal neurons

A growing body of literature suggests that hippocampal maps are not limited to representations of the physical environment; subsets of hippocampal neurons have been shown to “map” continuous scalar quantities other than physical space. In one study, rats ran on a treadmill to receive a reward (Kraus et al., 2013). Importantly, while running, the rat’s position was mostly stable. Rats were trained to run for a predictable amount of time (varied speed and distance), or predictable distance (varied speed and time). In this paradigm, a number of neurons that fired selectively for a specific time interval were found, while distance varied and physical location was fixed (Kraus et al., 2013). In another study, rodents were trained to hold a lever to play a tone that continuously increased in frequency, with the goal of releasing the lever within a learned frequency range (Aronov et al., 2017). Here, sound represented a non-spatial scalar quantity that could be continuously mapped. Indeed, hippocampal neurons showed tonal selectivity for all frequencies used in the experiment, and only when the tone had a predictable relationship with reward. Whether a neuron also showed a physical place specificity was not correlated with its tonal specificity (Aronov et al., 2017). These studies illustrate hippocampal ensemble-level encoding of non-spatial continuous sensory dimensions of an experience in rodents.

Clear evidence for pure spatial selectivity in navigating primates (analogous to that commonly observed in rodents) is scarce; studies that aim to disambiguate purely spatial coding from possible confounding factors describe primate hippocampal place cells as view-dependent, task-related, or otherwise complex (Ekstrom et al., 2003; Georges-François et al., 1999; Ono et al., 1993; Wirth et al., 2017). In previous work, the specific factors that modulate spatial activity in hippocampal neurons could not be disambiguated or contrasted with adaptive feature selectivity as shown in the current study; the use of single tasks with fixed object-or subject-positions makes it difficult to precisely ascribe this neuronal selectivity.

Most recently, primate hippocampal neurons have been ascribed gaze-informed, task-situated place selectivity (Wirth et al., 2017). However, interpretations of “action context” in this work rely on active versus passive movement through a restricted environment, rather than free navigation and environmental exploration in a common environment during varied cognitive tasks in the same space. In the current study, differential encoding of space across cognitive tasks can be attributed to specific selectivity for viewed and remembered objects that guide behaviour. These results extend the concept of mixed selectivity for cognitive variables seen in the primate prefrontal cortex neurons (Rigotti et al., 2013) to hippocampal representations of remembered features. Furthermore, single neurons with retrospective selectivity show that encoding of non-spatial features is not necessarily linked to incoming sensory inputs, but to memories of these features. These multidimensional memory fields provide a single flexible neural substrate for two main functions of the hippocampus: spatial navigation and associative memory.

### Reconciling spatial and non-spatial theories of hippocampal function

The hippocampus is believed to play an important role in navigation primarily because of the observation of place cells which encode the spatial position of the animal. These studies are typically performed in contrived, geometrical environments with experimenters monitoring few behavioral variables, such as the animal’s position or head direction. Hence, it is natural to focus on navigation and ignore the more general role of the hippocampus in encoding new structured memories. However, it is known that the hippocampus is not needed to navigate in familiar environments (see e.g. Clark, 2018; Redish, 1999), suggesting that it is important primarily in situations of navigation that require the formation of new memories. Recent theoretical works propose, similarly to what has been suggested by Eichenbaum (2017b), that the hippocampus is a memory system, which uses the statistics of the recent history to compress a stream of highly correlated sensory experiences. Sensory compression can be achieved using a simple network trained as a sparse auto-encoder (Gluck and Myers, 1993; Olshausen and Field, 1997). Recently, this type of network was used to compress sensory experiences of a simulated agent exploring a new environment, and was shown to naturally produce the spatial response properties of typical hippocampal neurons (Benna and Fusi, 2018). The compressed sensory representations improve the efficiency of information storage for memory, produce representations that can be highly biased towards relevant portions of the environment, and explain the elevated variability of the neuronal responses.

Other recent models of the hippocampus focus on the encoding of temporal sequences and assume that the hippocampus tries to compress memories by storing only the information that is relevant for predicting the next state of the environment (Hardcastle et al., 2017; Schiller et al., 2015; Stachenfeld et al., 2017) at the expense of the specific role it plays in navigation. These predictions are then used to drive reinforcement learning (Mattar and Daw, 2018; Stachenfeld et al., 2017). All these models are compatible with our observations and provide a general theoretical framework to understand the role of the hippocampus as a memory device, with spatial response fields that are modulated by cognitive factors and recent sensory experience.

### Conclusions

Instead of framing the hippocampus as the brain’s Global Positional System, the spatial, sensory, and mnemonic encoding observed in our study better reflect the processes inherent in Tulving’s General Abstract Processing System (Tulving, 1985). In such a system, adaptive representations provide the basis for learning and storing behaviorally relevant information across behaviorally relevant dimensions in a context-dependent manner.

## Acknowledgements

We thank Jesse Jackson, Matthew Leavitt, Ramon Noguiera for critical editing, input and discussion, and all members of the JMT lab for support. We thank Blandine Bally, Kevin Barker, Jackson Blonde, Sara Chisling, Steve Frey, Steve Nuara, and Walter Kucharski for technical assistance. RAG was supported by an NSERC PGS-D and a McGill David G. Guthrie Fellowship. This work was supported by CIHR and NSERC grants to JMT.

## Author Contributions

All authors edited the manuscript. RAG designed experiments, virtual environments, collected and analyzed data, and wrote the manuscript. LD contributed to data analysis. BWC contributed to data collection and data analysis. GD contributed to virtual environments design and data analysis. SW contributed to experimental design. SF contributed to data analysis and manuscript writing. JMT contributed to experimental design and manuscript writing.

## Methods

### EXPERIMENTAL MODEL AND SUBJECT DETAILS

Two male monkeys (*Macaca mulatta*; 7 years old, 7 kg; 14 years old, 12 kg) participated in all experiments. These monkeys were trained to perform three behavioral tasks, and given juice reward for their efforts in each task (400-1000+ mL daily). Monkeys also received food rewards as positive reinforcement at the beginning and end of each session. Behavioral patterns and body weights were closely monitored to ensure stable health conditions throughout the experiment. All animal procedures complied with the Canadian Council of Animal Care guidelines and were approved by the McGill University Animal Care Committee.

### METHOD DETAILS

#### Electrophysiological Recordings

The entire protocol for planning surgical procedures, recording from the hippocampus and verifying electrode locations is schematized in **Figure S1**.

Prior to any surgical procedures, a naïve 500um isotropic T1-weighted 3T MRI was taken for each animal (**Figure S1**, step 1). Using these scans, head post placement and chamber trajectory were planned using an MRI-guided neuro-navigation suite (Brainsight, Rogue Research, Montreal, QC) (**Figure S1**, step 2). Chambers were positioned over prefrontal cortex, such that electrode trajectories were perpendicular to the long and transverse axis of the right middle-to-posterior hippocampus. Following surgical implantation of the head post and recording chamber, a CT scan was acquired with cannulas passing through the chamber grid at cardinal locations (**Figure S1**, step 3). The resultant CT and MRI were co-registered, so electrode trajectories and terminal recording locations could be specifically mapped to chamber grid holes (**Figure S1**, step 4).

All data was collected over the course of 37 recording sessions. In each session, hippocampal activity was recorded using up to four single high-impedance tungsten electrodes (0.4-1.5 MOhms) simultaneously. Prior to every recording session, electrode trajectories were mapped to the MRI, and expected distances to grey and white matter were measured (**Figure S1**, step 5). These expected waypoints were compared against changes in neural activity while the electrode was lowered to the terminal recording site (speed 0.01mm/s; **Figure S1**, step 6). Distances to putative CA3 recording sites were adjusted online as necessary.

Neurons from the hippocampus were isolated while subjects sat quietly in the dark recording room, since hippocampal neurons typically exhibit elevated firing rates in this state compared to foraging or other exploratory behaviours. Local field potentials were monitored for bouts of theta-like activity and changing low frequency power profile as a function of arousal. Multi-unit activity was monitored for sparse activity and burstiness characteristic of hippocampal pyramidal neurons. Hippocampal activity was recorded at 30,000 Hz using a multi-channel recording system (128 channel Cerebus Data Acquisition System, Blackrock Microsystems, Utah, USA) for sorting and offline analysis. Cluster-cutting to isolate neurons from multi-unit clusters was done using Plexon software (Offline Sorter, Plexon Inc., Texas, USA). Cluster cutting was done agnostic to time; however, neurons with continuously morphing principle components and/or a complete loss of activity as a function of time were excluded from analyses. Any neurons with task-invariant reward-related activity were excluded from analyses.

In one monkey, post-recording verification of electrode trajectories was possible. This was done using a 350um isotropic susceptibility-weighted 7T MRI (**Figure S1**, step 7, cool color map). This scan was co-registered to the naïve 3T anatomical MRI, and shows a high degree of concordance between the expected and actual trajectories and terminal recording locations.

#### Behavioral Tasks

After electrode positions were fine-tuned to optimize neuronal signal-to-noise ratios, monkeys typically sat quietly in the darkened room for 20 minutes, allowing sufficient time for electrodes to settle. After this, monkeys proceeded to complete three behavioral tasks: a cued saccade task, a virtual associative memory task, and a virtual foraging task. At the end of each session, the cued saccade task was repeated, and monkeys again sat quietly in the dark for an additional 10-20 minutes of recording.

In the cued saccade task, monkeys were trained simply to fixate on a 1° visual angle (dva) white dot that could appear at any of 9 locations on the monitor in a 24 dva by 18 dva grid.

The remaining two tasks were incorporated into a virtual environment custom-built using a video game engine (Unreal Engine, Epic Games, Inc., Rockville, MD) and parameters of the behavioural tasks could be monitored and controlled in real-time via Matlab (Mathworks, Inc., Natick MA) (Doucet et al., 2016). Monkeys were trained to freely navigate about this environment using a two-axis joystick.

In the virtual foraging task, subjects were visually cued to navigate towards an easily identifiable target (red fog) that was consistently rewarded. The target could appear at any of 84 locations in the environment (**Figure 1C**, Foraging task, 84 dotted locations for display purposes only). Target locations in the X-Maze were independent across trials.

In the associative memory task, monkeys navigated the X-Maze to learn a context-dependent reward value hierarchy. Reward value associations were dependent on environmental cues (textures applied to maze walls; Context 1 and Context 2) and three differentially rewarded colored discs (objects A, B and C). Object reward values were context-dependent; that is, in Context 1, object A > B > C, and in Context 2, object C > B > A (**Figure 1C**, **Figure S3A**). Object colors were pseudo-randomly selected from a seven-color set at the beginning of each session to prevent repetition of colors across neighboring sessions. Thus, a new context-object hierarchy was learned every day.

On a single trial, subjects start at either the North or South end of the X-Maze. They then navigated through a long central corridor towards the opposite end of the maze. One of two possible textures (wood or steel) was applied to some walls of the maze when the monkey reached the corridor (**Figure S3B** and **C**, position a). When subjects reach the end of the corridor, they reach a forked decision point, with each of the two arms containing one of the three possible colored objects (**Figure S3B** and **C**, position b).

On individual trials, the context was independently randomized, as was the object color combination. Object colors were randomly assigned to either the left or right arm of the maze, and the same two colors could not appear in each arm of the maze on a single trial.

Quantities of juice reward given for successful completion were fixed between the cued saccade task, foraging task, and middle reward value of the associative memory task.

#### Experimental Set-Up

During training and experimental sessions, the monkeys were seated in a custom-built chair in front of a computer monitor. The chair was fit with a two-axis joystick (part 212S15S8383; PQ Controls, Inc., Bristol, CT), which the monkeys used to navigate freely through the virtual environment. Player position within the virtual environment was updated and recorded at 75 Hz, matching the monitor refresh rate. While seated, the head position of the monkey was fixed, to facilitate eye position and intra-hippocampal recordings. Eye position on the screen – and thus gaze within the virtual environment – was monitored at 500 Hz via video-based eye tracking (EyeLink 1000, SR Research, Ontario, Canada).

#### Eye movement classification

We use a custom toolbox to parse eye signal collected via video oculography into saccades, fixations, smooth pursuits and post-saccadic oscillations (Corrigan et al., 2017). Briefly, the initial identification of putative saccades is done by: 1) iteratively calculating a saccade acceleration threshold; 2) grouping threshold crossings within 40ms into a putative saccade; and 3) ignoring putative saccades group shorter than 10ms. The remaining segments of eye signal were further classified by foveation type.

For all putative saccadic periods, the maximum velocity was calculated, and then the onset and offset precisely identified by comparing the main direction and inter-sample changes in direction. Saccade boundaries were defined when the signal was either above a high threshold (60°) for one sample, or above a low threshold for three consecutive samples (20°). This method differentiates between eye movement types, since saccade direction is very consistent, whereas camera noise leads to higher inter-sample variance during smooth pursuits and fixations. Once saccades were identified, the direction and amplitude were calculated based on the onset and offset points for all saccades during the visually-guided task, including inter-trial-intervals, and for all completed trials in both virtual navigation tasks. These saccades were used for analyses of saccade direction selectivity. Saccade offset locations were used to analyze gaze position selectivity on the screen.

### QUANTIFICATION AND STATISTICAL ANALYSIS

#### Saccade Direction Selectivity

To examine saccade direction selectivity, we binned directions in eight 45° bins, starting with a center on 0°. A bin was only analyzed if there were at least 7 saccades in it, and a neuron’s saccade direction selectivity was only analyzed with a minimum of 5 saccade direction bins. Firing rate was calculated for the 150ms prior to saccade onset to get the average spike rate for each direction.

Significant direction selectivity for each neuron was assessed using a permutation test. Firing rates and directions were randomly shuffled 1000 times, generating 1000 null distributions for each saccade direction for each neuron. A neuron was categorized as being selective for a direction if it had a spike rate that was in the top 5th percentile of the null distribution after Bonferroni correction (alpha = 0.05/number of direction bins).

#### Gaze Position Selectivity

Each on-screen foveation was categorized within one of nine 12°×8° screen areas. For each foveation within each screen location, a neuron’s firing rate was calculated in the 200ms after saccade offset. Also for a location to be included, there needed to be at least 7 saccades per location. Foveation location selectivity was tested in each task for all neurons, with enough Foveations in at least 6 screen locations in each task. Like saccade direction selectivity, gaze position selectivity of each neuron was assessed using a permutation-derived null distribution.

#### Place Selectivity

To determine whether each neuron fired more action potentials than expected by chance in any area of the X-Maze in each task, we used a statistical permutation test based on spatial position and spike rasters for each neuron (**Figure 2**, **Figure 3**, and **Figure S4**). First, the X-Maze was parsed into a 32×12 grid of pixels. For each trial, a vector of player positions (occupied pixel number) was created at 1ms resolution. For each recorded neuron, a corresponding binary vector was produced with 1ms resolution, denoting the presence or absence of an action potential. By collapsing across trials, the occupied time and number of recorded action potentials was computed. To determine if this number was significantly above chance for each pixel, the vector of occupied pixels was circularly shifted for each trial, and the firing rate in each pixel was re-computed. This circular shuffling procedure was done 1000 times. Any pixel with an empirical firing rate exceeding percentile 1-alpha was statistically significant, where alpha=0.05/total number of occupied bins. Pixels with a total occupancy of less than 200ms were excluded. Place fields were defined as any of the 9 architecturally distinct maze areas (4 arms, 2 branches, 3 corridor sections; see **Figure 3** and **Figure 4A**, allocentric reference frame) with at least one statistically elevated spatial bin.

The specificity of each neuron’s spatial response map was quantified using spatial information content (Ravassard et al., 2013; Skaggs et al., 1993) (**Figure 3B** and **C**, **Figure S6**). Each neuron’s information content (*I*; in bits) is defined as

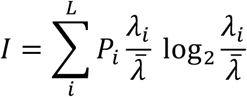

where the proportion of occupied time in each pixel (*P*_*i*_) is defined as

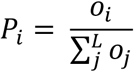

and the average firing rate per pixel 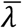 is

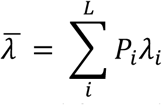

Spatial information content was computed for each neuron for each pixel occupied for more than 200ms. A null distribution of spatial information contents (and corresponding null spatial information maps) was computed for each neuron by circularly shifting the vector of occupied pixel numbers 1000 times prior to computing the spike rate maps for that neuron. Neurons with summed spatial information content exceeding the 95th percentile of the null distribution were deemed statistically significant, and these are included in the spatial histogram (Figure 3B), spatial information content histograms (Figure 3C, Figure S6) and cumulative distribution (Figure 3C, inset). Particular pixels with spatial information content exceeding that of their 95th percentile (Bonferroni-corrected for the number of occupied bins) were deemed statistically elevated, and displayed in the spatial histogram in Figure 3B.

#### Spatial Classification Analyses

We used a linear classifier (Fan et al., 2008) to determine whether the population of all recorded neurons could reliably encode position in the maze (**Figure 4**). The same procedures were used for all spatial classification analyses, starting with firing rates for all neurons in all trials and respective conditions (i.e. “trials” could refer to passes through each spatial bin or individual trial periods in the associative memory task). All neurons that did not have at least 10 trials in all conditions were excluded from our analyses (n=152 included). To begin, we randomly subsampled 10 trials from each condition for each neuron, creating an ensemble subsample. For subsequent classification, we used a linear kernel support vector machine, cross-validated with stratified k-folds. Specifically, we split the ensemble subsample into five stratified groups of trials (five folds), with four folds constituting the training set, and one fold reserved for testing. At this point, the firing rate of each neuron in all k-folds were z-scored using the mean and standard deviation of the training set only for that neuron (not the testing set). A linear kernel model was fit to the ensemble subsample using L1-regularized L2-loss support vector classification. An important benefit of using L1 regularization is automatic parameter selection on the model inputs; whereas L2 regularization yields parameter weights very close to zero, L1 regularization instead shunts weights directly to zero. This results in a trained model that is both sparse and more interpretable. The trained model was subsequently tested on the reserved testing fold to assess prediction accuracy. The procedure for normalization, model training and testing was repeated five times total, so each k-fold of the ensemble subsample was used as the testing set once. The entire procedure — starting from the 10-trial ensemble subsampling — was repeated 100 times, yielding a total of 500 iterations of the SVM testing procedure.

A permutation procedure was used to determine chance prediction accuracy in all cases. This proceeded similarly to the training and testing procedures described above. However, after creating the ensemble subsample and prior to splitting the ensemble subsample into stratified k-folds, the condition labels were randomly permuted, and classification analyses then proceeded exactly as previously described. This procedure was repeated 20 times for each ensemble subsample, yielding a total of 10,000 individual iterations of the SVM testing procedure.

It is important to note that each neuron’s firing rate was z-scored in the training set only and within each task independently prior to classification. This negates the possibility of spurious similarity of cross-task classification models attributed to within-neuron similarity in baseline firing rates across tasks, independent of area-specific changes in firing rate. Similarly, this negates the possibility of spurious dissimilarity of cross-task classification models attributed to within-neuron changes in baseline firing rates across tasks, independent of area-specific changes in firing rate.

#### Statistical evaluation of classifier performance

The mean and standard deviation classification accuracy reported herein include testing of each individual k-fold. Significant differences between accuracy distributions were tested using a two-sample Kolmogorov Smirnov test, Bonferroni-corrected for the total number of distribution comparisons.

Classification model reliability was further evaluated using Cohen’s kappa statistic (Cohen, 1960). The kappa statistic is an objective measure of classification reliability. Unlike raw or chance-normalized prediction accuracy, kappa provides a meaningful metric with which to compare the performance across classifiers — even with uncommon numbers of classes — because it is a bound statistic that relies on the observed and expected proportion of correct predictions for each class of a model; it is agnostic to the number of classes being differentiated. It is described as

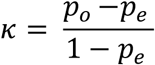

where *p*_*o*_ is the proportion of correct predictions, and *p*_*e*_ is the probability of guessing the correct class by chance.

#### Non-spatial feature selectivity

For two example neurons, we determined whether firing rate varied in each trial period as a function of non-spatial trial features (chosen object color, trial context, and their conjunction) from the current and previous trial (**Figure 5A** and **B**). Key trial events delineated trial periods. The post-reward period and pre-context period were equally split intervals of time between the start of the current trial (~200ms after reward from the previous trial ended) and the instant the subject reached the central corridor. The context appearance period spanned the entire length of the corridor, when the context was visible as a texture applied to the maze walls. Upon reaching the end of the corridor, the subject’s view in the maze was gently corrected to face cardinal direction north or south precisely, and subsequently both objects were triggered to appear simultaneously in the ends of the maze. This marked the start of the object appearance period. The object appearance period ended and the object approach period started ended on the same frame that the subject initiated a turn towards the chosen object for that trial.

We used multiple linear regression to determine whether each neuron’s firing rate was modulated as a function of non-spatial trial features (i.e., trial context, trial object colors, and their conjunction) in each trial period of the associative memory task. This procedure was repeated using trial features for the current and previous trial. Formally,

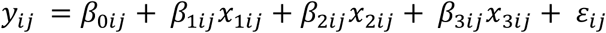

where *y* describes the change in a neuron’s firing rate within each task period (i; 1, post-reward period; 2, pre-context; 3, context appearance; 4, object appearance; 5, object approach) for current and previous trial features (*j*; 1, current trial features; 2, previous trial features). Fit parameter (*β*_0_) describes the intercept of the regression line. *e* estimates the residual. *β*_1_, *β*_2_, and *β*_3_ describe the effect of chosen object, trial context, and their conjunction, respectively. We assessed statistical significance of each of these parameters using a partial F-test, wherein the error of the full model is compared to that of a model with one parameter omitted. The proportion of all neurons (n=183) with significant fit parameters for object, context or their interaction is reported in **Figure 5C**.

#### Sensory versus Mnemonic Trial Feature Encoding

The F-statistic of the fit parameters *β*_1_, *β*_2_, and *β*_3_ were used to compare sensory and mnemonic encoding of associative memory trial features in individual neurons. Specifically, **Figure 7A** shows the scatterplot of neurons’ regression coefficient during the goal approach period (sensory encoding) versus the post-reward period (memory encoding).

The proportion of neurons with significant regression coefficients for each parameter was compared using McNemar’s test of proportions (**Figure 6A**, insets).

The relationship between sensory and memory encoding for each trial parameter for each neuron was characterized using the Pearson correlation coefficient rho (**Figure 6B**). Rho was calculated over 10,000 bootstrap iterations, and **Figure 6B** shows the median and 96.7% confidence interval (alpha = 0.05/number of trial parameters).

#### Trial Type Classification

Trial type classification (**Figure 7**) was done using Glmnet (Friedman et al., 2010) using firing rates from all neurons included in the spatial decoding analyses (n=152). For the Sensory condition, each neuron’s firing rate in each from the Corridor, Goal appearance and Goal approach periods were included as predictors. For the Memory condition, firing rates from the Post-reward and Pre-context periods were used as predictors. In the Sensory+Memory condition, all 5 periods were used. Because of the high ratio of model predictors to training and testing examples for these analyses, Glmnet classification was used with elastic net regularization. A nested cross-validation procedure was used to appropriately tune model hyperparameters (regularization parameter lambda, and elastic net L1-L2 weighting parameter alpha, via grid-search) and test on hold-out sets of trials never seen by the trained model.

## Supplementary Materials

### Online Movie 1. Foraging task trials

https://youtu.be/aWvheMzxMJo

Three example trials of the Foraging task from session W0325.

*Top:* LFP trace (white streaming line) and single unit action potentials (blue ticks, sound) recorded from unit W0325.A1M0.2.

*Circle and dot:* monkey’s eye position in the virtual environment.

*Bottom right:* time from trial start.

*Bottom left:* Name of every object that falls within 3 degrees of the foveated position.

### Online Movie 2. Associative memory task trials

https://youtu.be/RHx9Lw65oDw

Four example trials of the Foraging task from session W0325.

*Top:* LFP trace (white streaming line) and single unit action potentials (blue ticks, sound) recorded from unit W0325.A1M0.2.

*Circle and dot:* monkey’s eye position in the virtual environment.

*Bottom right:* time from trial start.

*Bottom left:* Name of every object that falls within 3 degrees of the foveated position.

---

**Figure S1.**
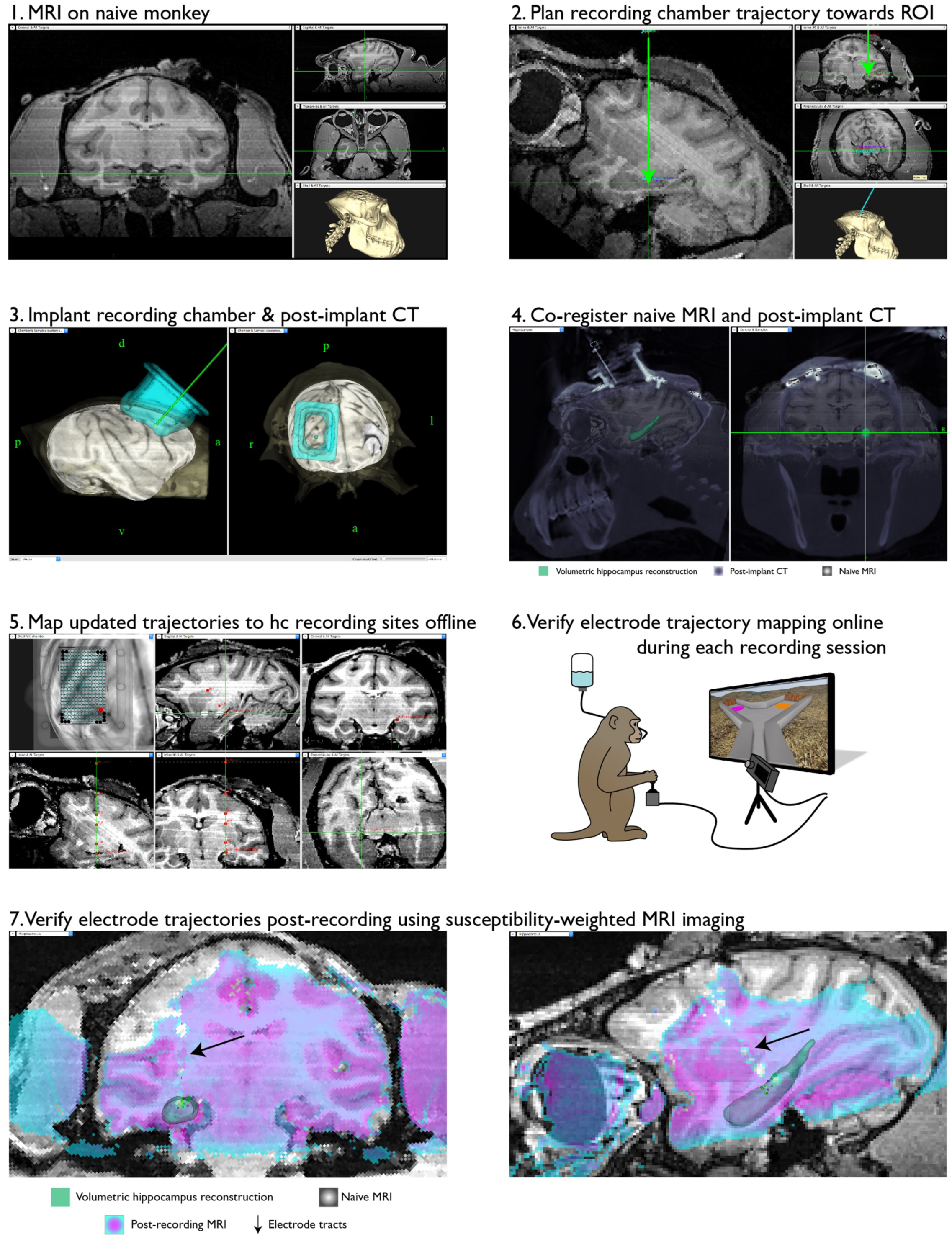
Hippocampal recordings: planning, mapping and verification. *Related to Figure 1*. Schematic representation of the major steps in planning, mapping, and verification of electrode trajectories and recording sites.

**Figure S2.**
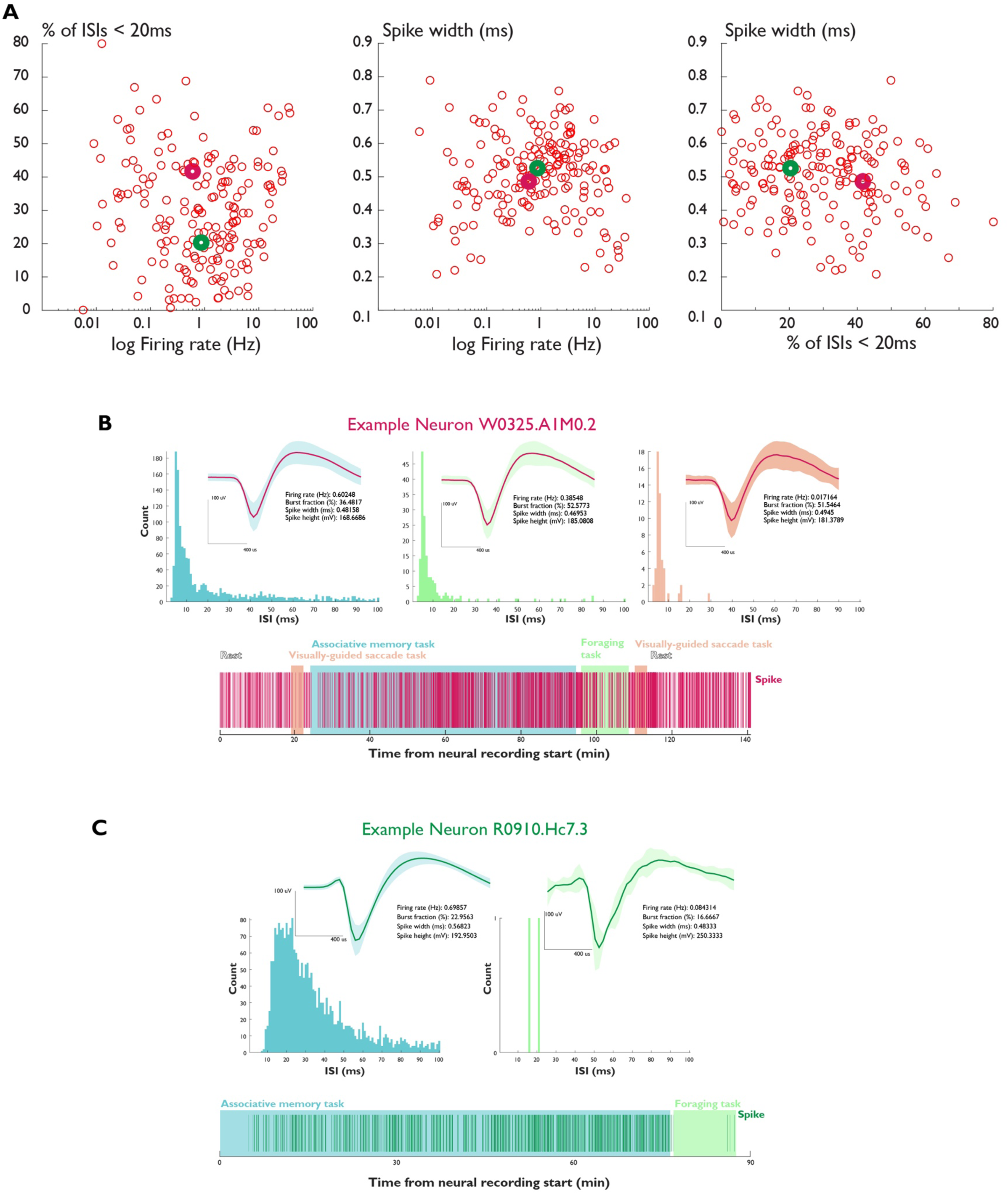
Individual neuron characteristics and example neurons. *Related to Figure 1*. (A) Burst fraction, spike width, and firing rate of all recorded neurons (n = 183). Purple and green circles mark the example neurons shown in (B) and (C), respectively. (B) Example neuron W0325.A1M0.2 inter-spike-interval distribution and average waveform. Shaded area, SEM. (C) Example neuron R0910.Hc7.3 inter-spike-interval distribution and average waveform. Shaded area, SEM.

**Figure S3.**
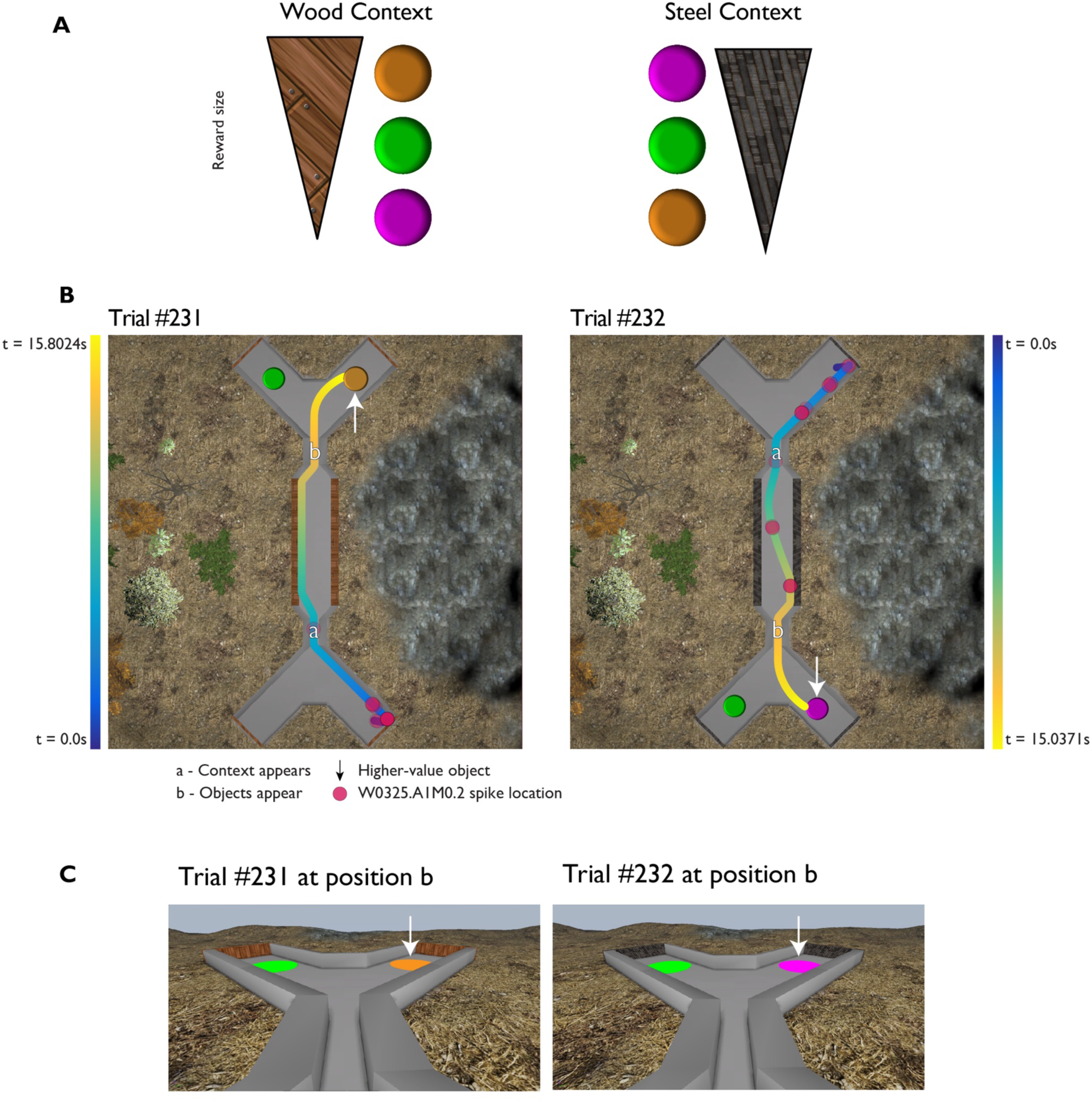
Example associative memory task reward hierarchy. *Related to Figure 1.* (A) Example of the reversed two-context, three-object reward value hierarchy for recording session W0325. (B) Two example trials from the recording session. Subject trajectories through the maze are colored according to the time from trial start (color bar). White arrow indicates the object of higher reward value. (C) Approximate first-person-view of the monkeys during each trial at position b. White arrow indicates the object of higher reward value.

**Figure S4.**
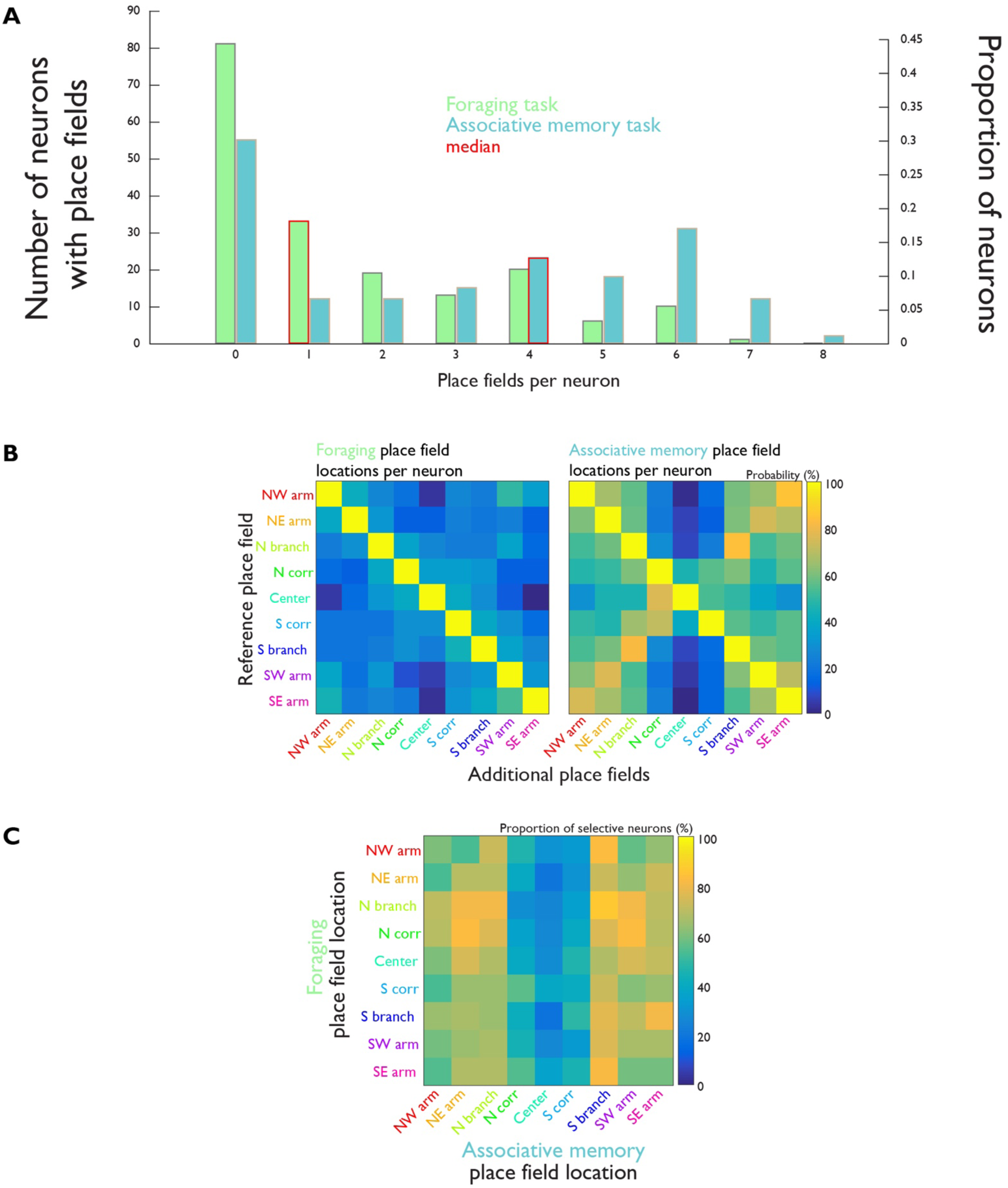
Single neuron place fields in each task. *Related to Figure 3*. (A) Number of place fields per neuron in each task. (B) Locations of concident place fields for all neurons with more than one place field in each task. (C) Location of coincident place fields for all neurons with at least one place field in each task.

**Figure S5.**
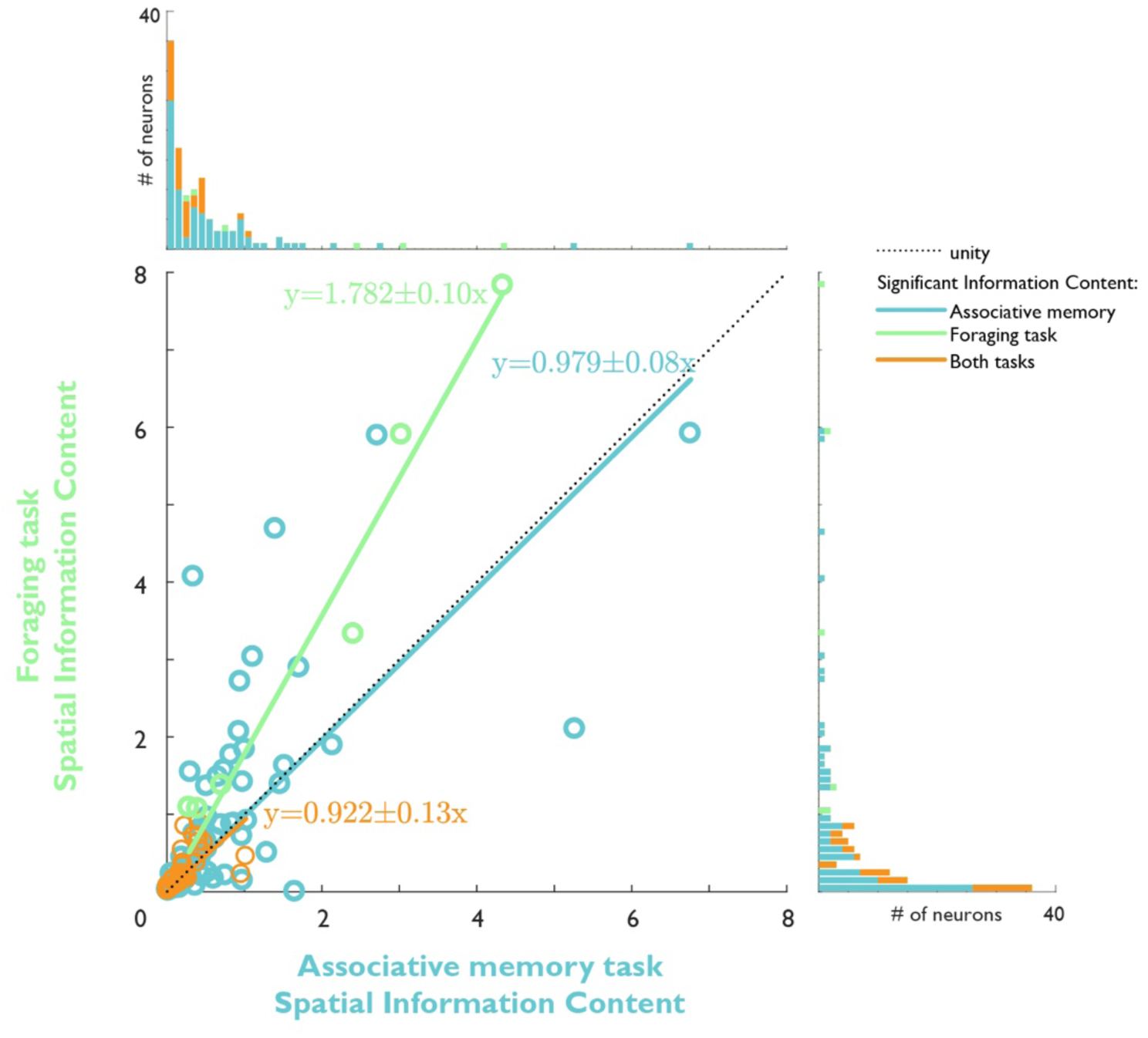
Comparison of significant neurons’ spatial information content in each task. *Related to Figure 3*.

**Figure S6.**
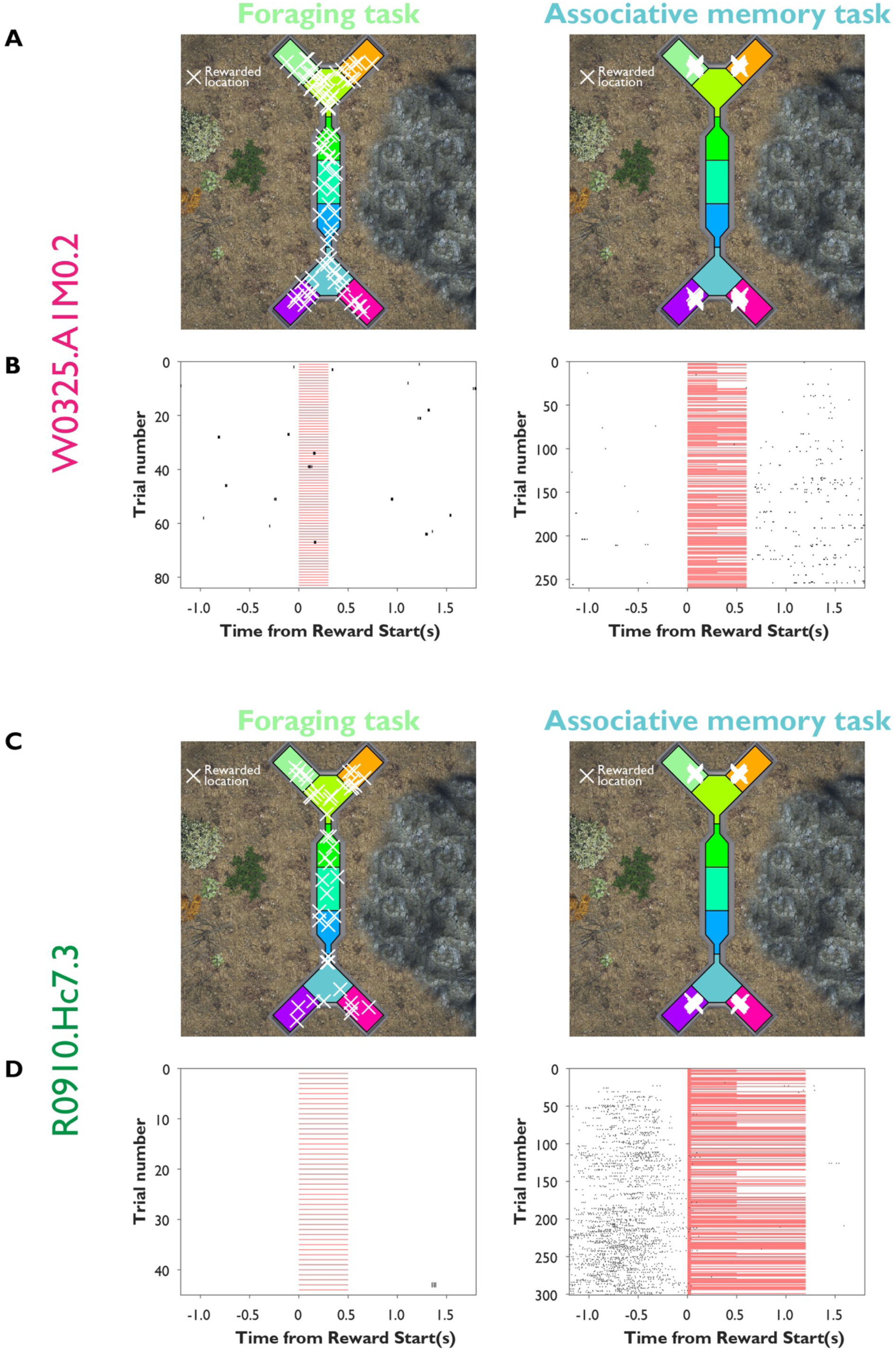
Rewarded-aligned spike rasters. *Related to Figure 4*. (A) Rewarded locations in session W0325 during the Foraging task (left) and Associative memory task (right). (B) Reward-aligned rasters for example neuron W0325.A1M0.2 in each task. (C) Rewarded locations in session R0910 during the Foraging task (left) and Associative memory task (right). (D) Reward-aligned rasters for example neuron R0910.Hc7.3 in each task.

